# Inference and Uncertainty Quantification of Stochastic Gene Expression via Synthetic Models

**DOI:** 10.1101/2022.01.25.477666

**Authors:** Kaan Öcal, Michael U. Gutmann, Guido Sanguinetti, Ramon Grima

## Abstract

Estimating uncertainty in model predictions is a central task in quantitative biology. Biological models at the single-cell level are intrinsically stochastic and nonlinear, creating formidable challenges for their statistical estimation which inevitably has to rely on approximations that trade accuracy for tractability. Despite intensive interest, a sweet spot in this trade off has not been found yet. We propose a flexible procedure for uncertainty quantification in a wide class of reaction networks describing stochastic gene expression including those with feedback. The method is based on creating a tractable coarse-graining of the model that is learned from simulations, a *synthetic model*, to approximate the likelihood function. We demonstrate that synthetic models can substantially outperform state-of-the-art approaches on a number of nontrivial systems and datasets, yielding an accurate and computationally viable solution to uncertainty quantification in stochastic models of gene expression.

## 1 Introduction

The stochasticity of biological processes at the single-cell level is one of the major paradigm shifts of twenty-first century biology [1–3]. Modern experimental methods, ranging from advanced microscopy to single-cell sequencing [4–6], have confirmed and detailed the pervasiveness of stochasticity in cellular biology. While these discoveries open new perspectives on the fundamental functioning of living systems, they also create novel challenges towards the development of mathematical models of biological processes, accentuating the role of statistical inference and uncertainty quantification in any modelling effort.

Biological variability is the result of many concomitant processes. A major source of noise (intrinsic noise) stems from the random timing of chemical reactions and is particularly important for reaction systems involving a small number of molecules of a certain species, as in many gene regulatory systems. The Chemical Master Equation (CME) [7] has been broadly adopted as a general framework to describe the intrinsic stochastic dynamics of chemical reaction networks [8]. While the CME benefits from an elegant mathematical formulation, its exact analytical solution is only known in a few instances [8]; on the other hand, the Stochastic Simulation Algorithm (SSA) [9] provides a Monte Carlo method to perform simulations of systems described by the CME.

Bayesian inference, the gold standard for capturing model and parameter uncertainty, relies on the likelihood function *p*(**x**_obs_ | ***θ***) to estimate parameters ***θ*** given observations **x**_obs_. For biochemical reaction networks, computing the likelihood requires a closed form expression for the solution of the CME, which is generally unavailable: while the forward problem of generating samples from the CME can be solved efficiently using the SSA, the backward problem of computing the probability of samples cannot. As a consequence Bayesian inference for biochemical reaction networks often relies on a variety of approximations to the likelihood function [8, 10].

Among the most well-known of these are the Finite State Projection (FSP) [11], continuum approximations [7, 12], and moment equations [13]. The FSP solves the CME on a finite truncation of the state space, whose size typically grows exponentially in the number of species; in practice this approach relies on computationally intensive approximations [14–16] for more complex systems. Continuum approximations to the CME based on stochastic differential equations, such as the Chemical Langevin formalism [12] and the Linear Noise Approximation [7] are limited to systems with small noise and in the case of the latter, Gaussian copy number distributions. Moment equations can be derived from the CME and used to construct an approximate likelihood function [17]; for systems with bimolecular reactions the moment equations have to be “closed” by a process called moment-closure, which yields approximate solutions of highly variable quality [13]. These approaches, termed Moment-Based Inference (MBI), are commonly used in practice [17–23] and usually very scalable, but the error introduced by the approximations can be difficult to quantify. Recent work [23–25] has pointed out that these methods can perform poorly for some systems, leading to biased or overly uncertain parameter estimates.

An alternative to analytical approximations of the CME is provided by *simulator-based* inference [26], which relies on simulations of the original model to estimate the likelihood using Monte Carlo methods. This family of methods only requires the ability to perform simulations of the model, which for biochemical reaction networks can be readily obtained using the SSA. Simulator-based approaches are mostly model-agnostic and can be easily adapted to many different problems, but due to their generality they typically require many simulations to produce a fully data-driven approximation of the likelihood.

Perhaps the best-known simulator-based inference method is Approximate Bayesian Computation (ABC) [27, 28]. ABC replaces the likelihood *p*(**x**_obs_|***θ***) with *p*(*d*(**x**_obs_, **x**) ≤ *ϵ* | ***θ***), the probability that the model generates outputs within a tolerance *ϵ* of the observed data, where *d*(·,·) denotes an appropriately chosen discrepancy measure. The posterior is then estimated by repeatedly sampling parameters and accepting those falling within this threshold. Tuning the discrepancy measure and the parameter *ϵ*, which trades accuracy for number of simulations, is difficult in practice and usually requires compressing the model output into low-dimensional summary statistics, a step that typically entails a loss of information.

A different simulator-based approach is Synthetic Likelihoods [29, 30], where the likelihood is approximated by a multivariate Gaussian whose mean and covariance are estimated from simulations. We will refer to this method as Gaussian Synthetic Likelihoods (GSL). Like ABC this approach frequently works with summary statistics of the data, which in this case should be approximately normally distributed under the model. For population snapshot and time-series data obtained from biochemical reaction networks GSL is unlikely to provide accurate results [31, 32]: indeed, analytically derived Gaussian approximations to the CME are commonly used in inference [19, 22, 33], but as shown in [24] these can result in unusable parameter estimates for some systems. Methods that approximate the likelihood based on kernel density estimation [34] or neural networks [35] can bypass this limitation, but they can require significant amounts of tuning and computational power to work well. A scalable approach to inference would ideally combine the flexibility of simulator-based methods with prior knowledge of the model to provide efficient yet flexible means of approximating the likelihood function.

In this paper we propose a new method for inference in a wide class of biochemical reaction networks, specifically those modelling gene expression, which is rooted in the specific characteristics exhibited by models of gene regulatory networks. Gene expression systems can often be thought of systems switching between discrete states of expression, broadly speaking corresponding to patterns of activation states of the genes’ promoters [36–39]. It is therefore natural to abstract the dynamics of gene systems as an indirectly observed dynamical system over a discrete (finite) set of states. These states are measured through observations of molecular counts; motivated by experimental measurements of the distributions of transcript and protein numbers, as well as analytical solutions of the CME obtained in a variety of cases, we propose a negative binomial mixture distribution as a model for molecular counts in our coarse-grained models of gene expression. We therefore propose that mixtures of time-dependent negative binomials can provide a tractable class of approximate models of gene expression systems. We call this class of models *synthetic models* (SM), and use these to approximate the likelihood function of gene expression models by fitting them to model simulations, in the spirit of Synthetic Likelihoods [30]. In cases when measurements are taken at short time intervals, where time correlations are particularly important, SMs can be further enhanced by imposing Hidden Markov Model dynamics on the latent states. We show that SMs can provide excellent estimates of the model likelihood where other methods (FSP, GSL, MBI and ABC) struggle, and that our approach can be applied to obtain accurate parameter and uncertainty estimates for challenging inference problems.

## 2 Synthetic Models

Our approach to inference is based on approximating the distributions predicted by the CME within a suitable family of candidates. We are in particular interested in gene expression systems, including those with feedback, but our methodology is general and can be applied to a large class of models, including non-Markovian models such as those recently considered in [40, 41]. An outline summarizing the method can be found in Fig. 1**a** and **b**.

**Figure 1:**
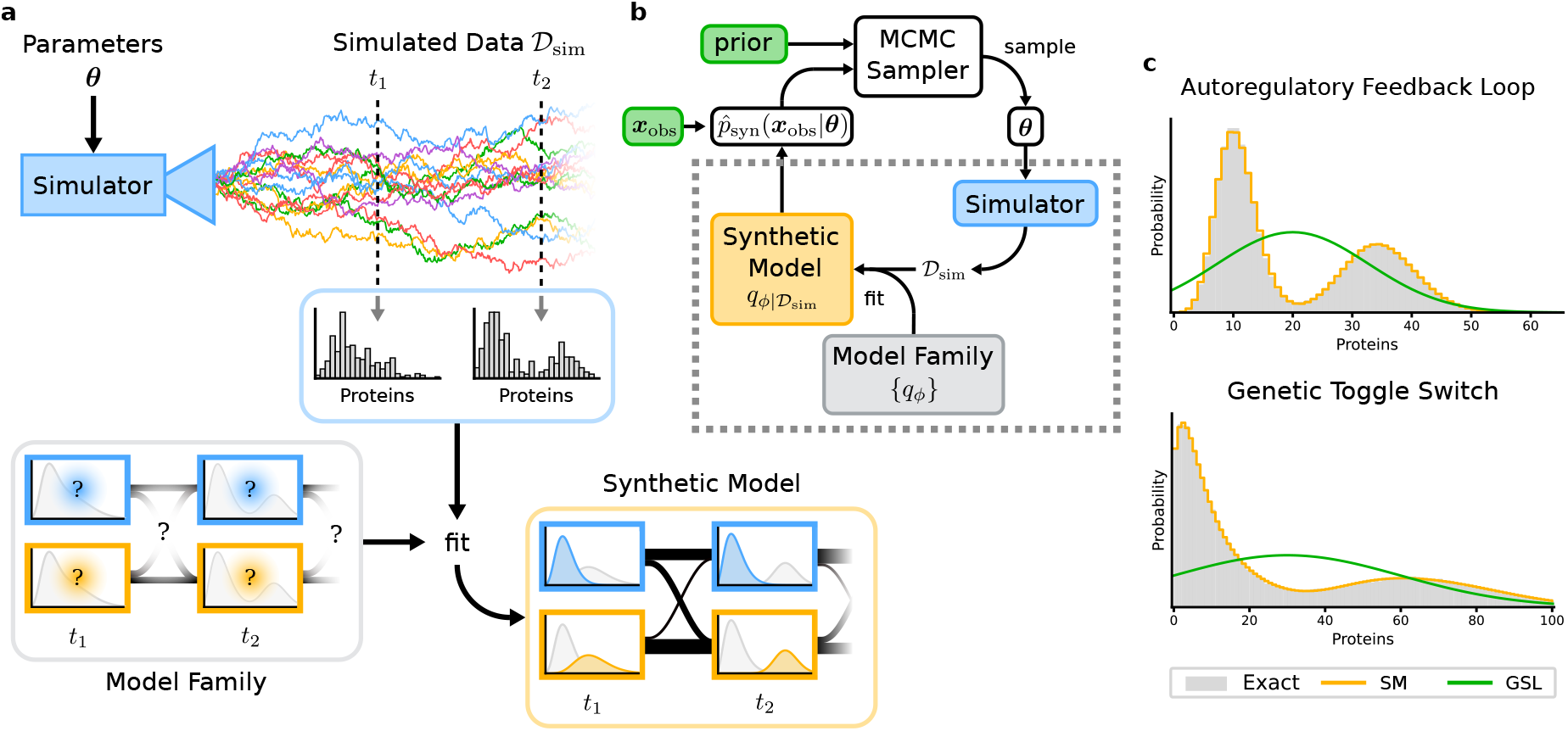
Bayesian inference using synthetic models. **a**: Given a parameter set ***θ***, stochastic simulations of the original model are run to obtain samples 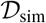. The synthetic model is picked from a parametric family of candidates and fit to the simulated samples. Our synthetic model is a finite-state Markov model with negative binomial output distributions for each state. **b**: Synthetic models (dashed box) provide an approximation to the likelihood 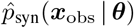 that can be plugged into standard MCMC algorithms for parameter inference and Bayesian model selection. **c**: Synthetic models based on negative binomial output distributions typically provide better fits to data than Gaussian approximations

Theoretical investigations have shown that single-time marginal distributions predicted by the CME for a variety of models describing many of the major biomolecular processes affecting gene expression (transcription, translation, cell growth, DNA replication and cell division) can be approximated by mixtures of negative binomials (MNBs) in the presence of timescale separation [42–46] – for an illustration see Fig. 1**c**. When timescale separation is not applicable, such an approximation cannot be derived analytically, yet measurements of the distribution of mRNA and protein numbers in bacterial, yeast and mammalian cells show that these are still well fit by such mixtures in many cases [36–39].

Having established that mixtures of negative binomials provide a good statistical model for experimental measurements of gene expression networks at fixed times, the next step is to extend this to include time dependency. Experimental measurements of a system at different times will be correlated, and a natural way to emulate these correlations is to treat the individual mixture components at each time point as states in a Hidden Markov Model (HMM). More precisely, we propose using a finite-state Markov chain with negative binomial output distributions for each state, see Fig. 1**a** for an illustration. This statistically tractable surrogate model, which we term synthetic model, defines a surrogate distribution over observations jointly at all measured time points. Note that integrating out the hidden state variable shows that the marginal distribution at any time point is still a mixture of negative binomials.

Assuming the true distribution *p*(**x** | ***θ***, *t*) predicted by the CME can be approximated by a mixture of negative binomials *q_ϕ_*(**x**) with parameters *ϕ*, a principled way to determine these parameters is to minimise the Kullback-Leibler divergence between the two distributions:

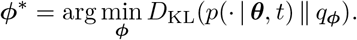

Here *q_ϕ_* is the MNB with mixture parameters *ϕ*. Since the reference distribution, being given by the solution of the CME is in general inaccessible, we can approximate it empirically by drawing samples using the SSA; minimising the above KL divergence is then equivalent to maximising the likelihood of the simulated samples, up to sampling error.

Fitting mixtures of negative binomials to data can be done efficiently using the Expectation-Maximisation (EM) algorithm described in [47, 48]. In order to fit all parameters of a HMM, including the initial distribution and the transition rates, we used the Baum-Welch algorithm (see SI for details), a type of EM algorithm applicable to HMMS. Once we have fit our synthetic model we can then compute the likelihood of our experimental observations **x**_obs_ using the forward algorithm for HMMS. This likelihood can be used to compute the posterior over parameters ***θ***, typically using MCMC, or to find the most likely parameters via optimisation – for an illustration see Fig. 1**b**.

Our procedure to estimate the likelihood *p*(**x**_obs_ | ***θ***) for model parameters ***θ*** is as follows:

1. Simulate sample trajectories using the SSA for the original model with parameters ***θ***
2. Fit the parameters of the HMM to the simulated trajectories using the Baum-Welch algorithm
3. Evaluate the fitted model at the observed data **x**_obs_ to obtain the synthetic likelihood 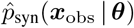.

Note that the synthetic model has to be fit from scratch for every parameter set at which the likelihood is queried, which is the main computational bottleneck of our approach.

The number of simulations and mixture components should be chosen appropriately for the reaction network. In our experiments we simulated each system at several randomly chosen parameters and ensured that the given number of simulations and mixture components could accurately reproduce the observed distributions. In an MCMC context, the number of simulations should be chosen such that the variance of the likelihood estimate likelihood still results in an acceptable rejection rate. Allowing a few more components than necessary did not affect the quality of fit in our experiments, as extraneous components either merged with others or were assigned negligible weights.

We remark that experimental data for mRNA or protein number generally comes in the form of either population snapshot data or live cell imaging. In the case of the former, each snapshot represents a different group of cells and modelling correlations at different times becomes unnecessary; it therefore suffices to fit mixtures of negative binomials independently for each time at which a snapshot is taken. This simplification can also be made when time correlations are weak enough to be neglected, as is the case for the toggle switch model considered in the next section.

## 3 Results

### 3.1 Autoregulatory Genetic Feedback Loop

We consider an autoregulatory genetic feedback loop that is illustrated in Fig. 2**a**. It consists of a gene with two promoter states G_u_ and G_b_, and a protein P that is produced at different rates *ρ_u_* and *ρ_b_* depending on the promoter state. Protein production occurs in geometrically distributed bursts with mean burst size *b*. The promoter switches from state G_u_ to G_b_ by binding a protein molecule with rate *σ_b_*, and this process is reversible with rate *σ_u_*. Protein dilution is effectively modelled by a first-order reaction; note that all other rates are rescaled by the protein dilution rate. We assume mass action kinetics for all reactions. This is the prototypical example of stochastic self-regulation in a gene and can be rigorously derived from a more detailed model incorporating mRNA dynamics [46].

**Figure 2:**
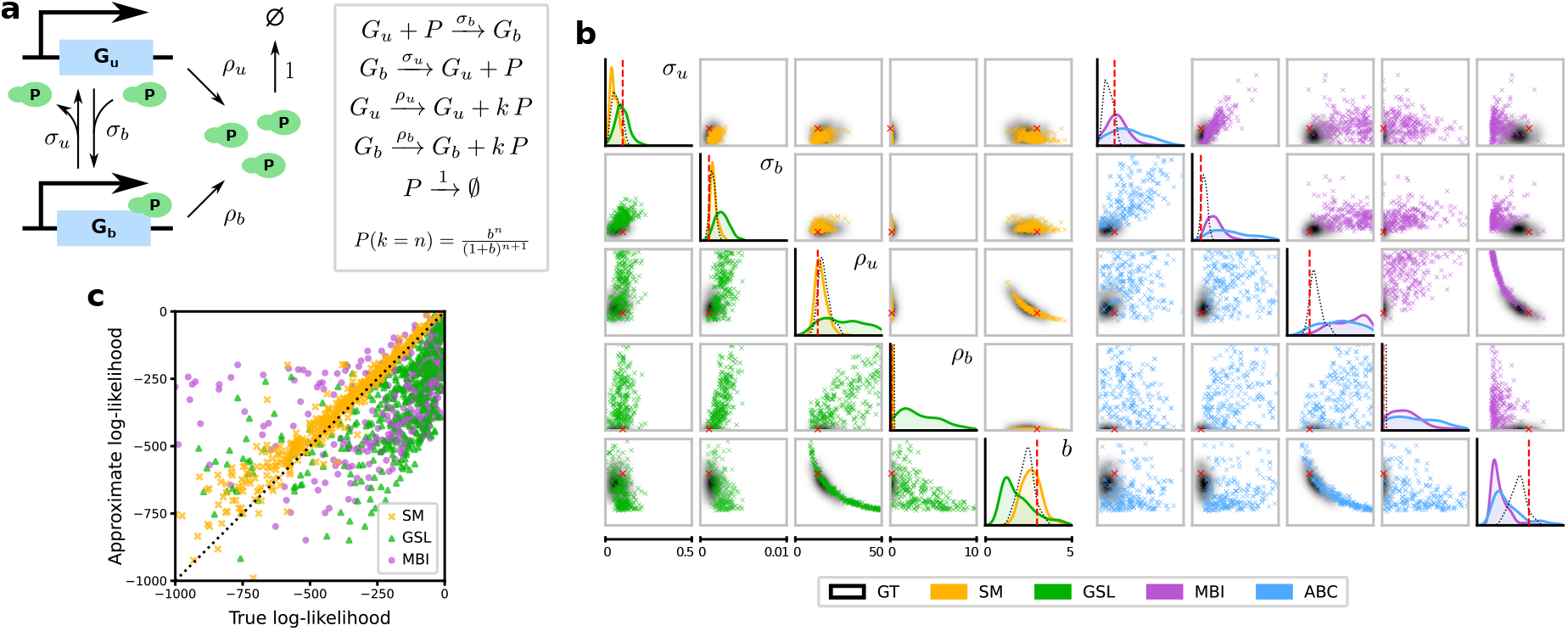
Comparison of Synthetic Models with standard methods for the case of an autoregulatory genetic negative feedback loop. **a**: Illustration of the reaction scheme describing the genetic circuit. **b**: Posteriors obtained using four different inference methods, with the ground truth solution computed using the FSP (black). The red dashed lines show the true parameter values. *Left*: Synthetic models (SM) and Gaussian synthetic likelihoods (GSL). *Right*: Moment-based inference (MBI) and sequential ABC. The ranges plotted coincide with the prior ranges. **c**: Comparison of true and approximate log-likelihoods. Parameter values were sampled from the prior and the true log-likelihoods were computed using the FSP. Synthetic models (yellow) provide significantly closer approximations to the true log-likelihood than either Gaussian synthetic likelihoods (green) or moment-based likelihoods (purple). The true parameter values are given in SI Fig. S1. The input data consists of protein numbers from 25 SSA trajectories measured at times *t* = 4, 8, 12, 16.

We consider the negative feedback regime where the protein production rate decreases upon protein binding (i.e. *ρ_b_* < *ρ_u_*). Due to the simplicity of this model, likelihoods can be efficiently computed using the FSP using a truncation to several hundred states in our examples, leading to an essentially exact solution and enabling us to compare our method with exact Bayesian inference. We tested our approach by using the SSA to simulate time-series data from several genetically identical cells and performed Bayesian inference based on the observed protein numbers, with a uniform box prior on the model parameters. For all methods except ABC we sampled from the posterior using the Metropolis-Hastings sampler with a fixed Gaussian transition kernel (see SI Section 1 for details). We note that while the steady-state solution of the CME of this system is predicted by theory to be well approximated by a negative binomial mixture (because of the small promoter switching rates compared to other rates [46]), we use data collected in pre-steady state where theoretical results are difficult to obtain. Hence the use of SM as a means to automatically obtain a negative binomial mixture approximation of the likelihood is particularly useful in this case.

Due to the presence of bimolecular protein-gene interactions, solving the moment equations for moment-based inference in this model requires a moment-closure approximation. We used the Linear Mapping Approximation (LMA) [49] for this purpose, which provided very accurate moment estimates for the parameter ranges considered in our experiments.

We compared the exact posterior obtained using the FSP with those computed using synthetic models and three representative inference methods: GSL, MBI and ABC (Fig. 2**b**) – see Methods and SI for details. For all parameters, the mode of the posterior computed using FSP or Synthetic Models is close to the true parameter values; this is not the case for the other methods. In particular our approach was the only one to yield a posterior where *ρ_b_* was concentrated around the true value of zero, whereas the other methods yielded posterior means that were significantly nonzero, falsely suggesting leaky gene expression. This example illustrates that using detailed distributional information can be valuable for discriminating between different modes of gene expression. Fig. 2**c** shows that synthetic models approximate the true likelihood of the model substantially better than both GSL and MBI, uniformly over the range of parameters considered (ABC does not yield explicit likelihood estimates). See SI Fig. S2 for further data including the posterior and MLE predictive distributions obtained using these methods.

In SI Fig. S2 and S3, we repeat the same analysis for a positive feedback loop where the protein production rate increases upon protein binding (*ρ_b_* > *ρ_u_*). As for the negative feeback loop, we find that likelihood approximation and parameter inference using synthetic models is significantly more accurate than using standard methods.

### 3.2 Genetic Toggle Switch

Next we consider a genetic toggle switch [50] in a eukaryotic cell (for an illustration see Fig. 3**a**). This consists of two different promoters, each of which can be on or off, and the protein from each promoter represses the expression of the other. We explicity model the translocation of mRNAs from the nucleus to the cytoplasm, the translation of cytoplasmic mRNAs into proteins and the translocation of proteins to the nucleus.

**Figure 3:**
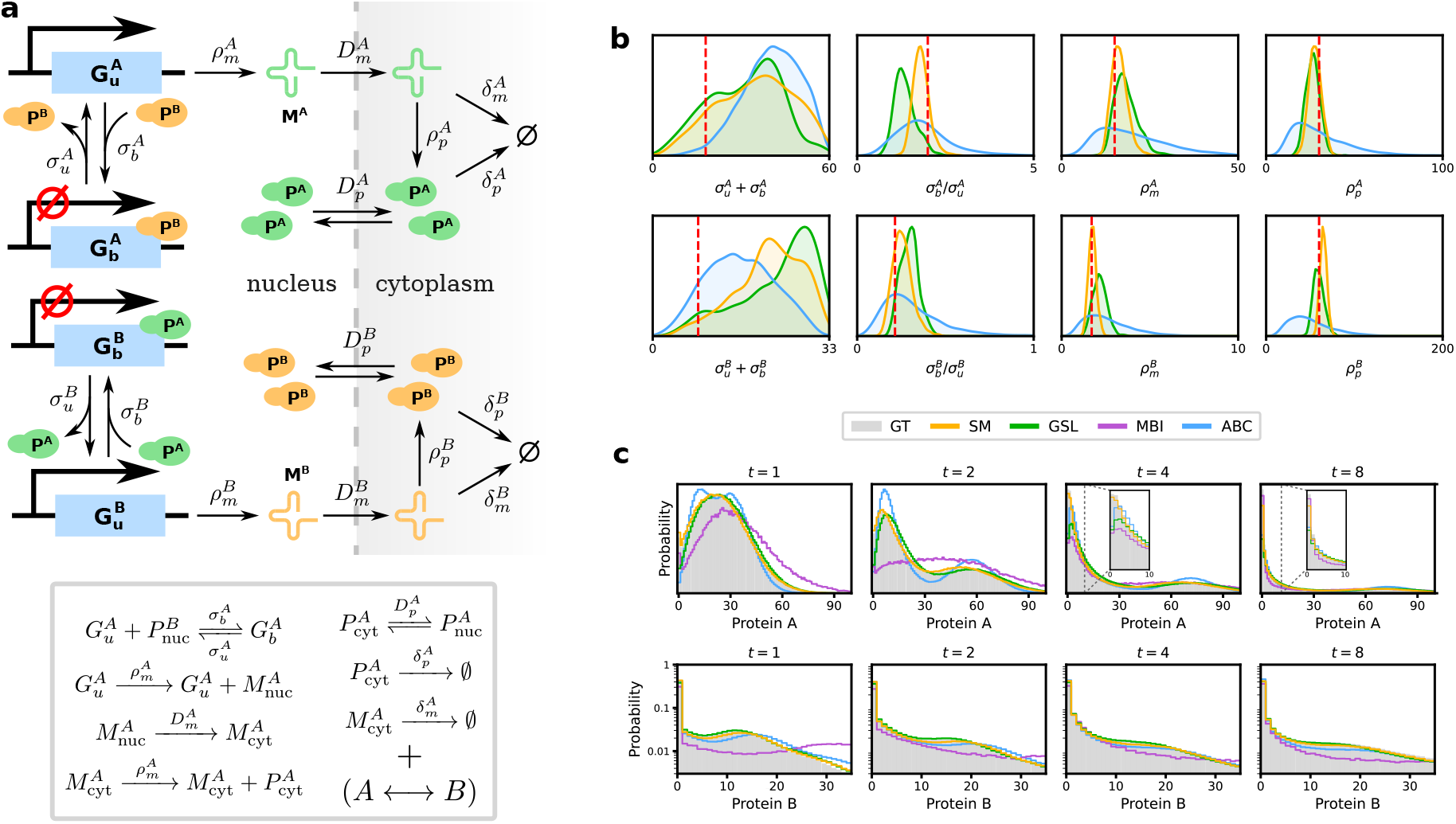
Comparison of Synthetic Models with standard methods for the case of a genetic toggle switch in a eukaryotic cell. **a**: Illustration of the reaction scheme describing the circuit, which is symmetric in *A* and *B.* **b**: Posteriors obtained using Synthetic models, Gaussian synthetic likelihoods, moment-based inference and Sequential ABC. **c**: Predictive distributions generated using the SSA at 4 time points for the maximum likelihood parameters obtained using each method. Synthetic models (yellow) and Gaussian synthetic likelihoods (green) provide significantly closer approximations to the true distribution (grey) than the other methods, with the former yielding a consistently more accurate fit for low molecule numbers (see insets). Parameters and prior ranges for all parameters are given in SI Fig. S4. The input data consists of cytoplasmic protein numbers (A and B) from 100 SSA trajectories measured at times *t* = 1, 2,…, 8.

This system is significantly more complex than the autoregulatory feedback loop considered above, involving an effective 10 species (we do not count bound promoter states due to conservation laws). For realistic mRNA and protein abundances (few tens and several tens to hundreds, respectively) a simple state space truncation would need to consider on the order of 10^9^ states, many orders of magnitude more than the previous example. Due to hardware constraints we were therefore not able to apply the FSP to this example; this illustrates the lack of scalability of the direct approach when applied to more realistic systems and the need for more efficient methods.

Fixing the translocation and degradation rates, which can often be deduced experimentally, we tested our approach in this case by inferring the remaining 8 parameter values. We used the SSA to simulate a synthetic dataset of 100 cells observed at eight different time points each, and performed Bayesian inference on the cytoplasmic protein numbers (both species) with a box prior around the true parameters (Fig. 3**b**) using Synthetic Models, Gaussian Synthetic Likelihoods, Moment-Based Inference (not shown, see below) and ABC. As with the autoregulatory feedback loop we used a Metropolis-Hastings sampler with a Gaussian transition kernel for all methods except ABC (see SI for details). Not all parameters of this model were identifiable from the data: while the ratio between the binding and unbinding rates for each gene can be identified, the individual rates themselves cannot. These findings did not depend on the method used, which suggests that we are dealing with intrinsic, as opposed to technical, unidentifiability. This is supported by SI Fig. S4, which shows that the predictive uncertainty in the posteriors is very small despite large variations in these two quantities. In contrast, the peaked posteriors around the true values of the transcription and translation rates show that these rates can well estimated by SM and GSL (which is not the case for ABC and MBI). We furthermore compared the predictive distributions for the maximum likelihood parameters estimated during inference (Fig. 3**c**) – we note that the SM prediction is the only one of all methods that is accurate for all times.

While the input to this experiment consisted of time-series data for SM, fitting a full HMM performed similarly to fitting independent mixtures of negative binomials at each time point, and we therefore used the latter approach for simplicity. We observed that using a full HMM for this model was more prone to local optima during the fitting step, which resulted in a higher variance of the approximate likelihood and reduced acceptance rates. Gaussian synthetic likelihoods similarly had significantly lower acceptance rates compared to independent mixtures of negative binomials, with a correspondingly increased number of MCMC iterations until convergence.

As for the autoregulatory feedback due to the nonlinearity of the propensities of the protein-gene interactions, moment-based inference for this model requires a moment-closure approximation. Out of the 9 different schemes implemented in the package `MomentClosure.jl` [51], the Linear Mapping Approximation (LMA) [23, 49] was the only one that consistently predicted positive moments around the true parameters, a necessary condition to get well-defined likelihoods. However even the LMA failed to predict the moments accurately for this system, resulting in a wildly skewed posterior (not shown) and heavily divergent predictive distribution (Fig 3**c**).

### 3.3 MAPK Pathway in *S. Cerevisiae*

Our final example uses experimental data from [52] to analyse the high osmolarity glycerol (HOG) MAPK pathway in *S. Cerevisiae*, where population snapshots were taken at different times after the induction of osmotic shock. The model is described in Fig. 4**a** which features highly non-Gaussian distributions of mRNA copy numbers. It consists of a single gene (STL1) in four possible states, each of which produces mRNA at a specified rate. Switching into one specified state is controlled by a kinase that is activated by a signalling cascade under osmotic shock; the concentration of the kinase is given as an external input to the system.

**Figure 4:**
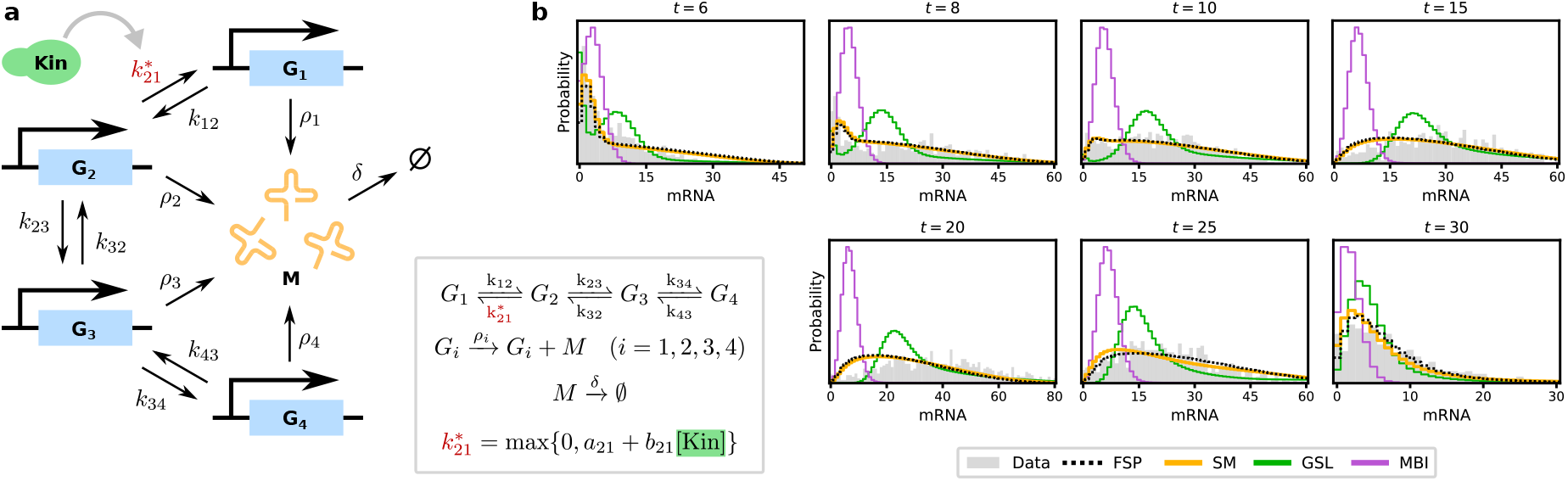
Comparison of Synthetic Models with standard methods for the MAPK pathway model. **a**: Illustration of the reaction scheme of the model, which consists of a gene in four possible states *G_i_* and mRNA. A kinase, whose concentration is a time-dependent input signal, modulates the transition rate 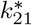 (see SI for details). **b**: Comparison of the experimentally observed distribution (grey) with the predictive distributions for the maximum likelihood estimates obtained using four different methods. Synthetic models (yellow) provide a quality of fit similar to the Finite State Projection (dotted line), whereas Gaussian synthetic likelihoods (green) and moment-based inference (purple) fail to capture the long-tailed shape of the distributions. Estimated parameters for each method are given in SI Table S1.

It was found in [24] that moment-based inference generally fails to yield good predictive results for this example, in contrast to direct likelihood-based inference using the FSP. As the model contains only 4 effective species (mRNA and 3 gene states, the fourth being given by the conservation law) it is very amenable to the FSP as a truncation to a few hundred states suffices to capture its dynamics in the relevant parameter range. Despite this simplicity, the model has 12 free parameters and poses a challenge for full Bayesian inference. A random-walk Metropolis-Hastings algorithm would require very small step sizes in order to keep acceptance rates high in 12 dimensions, requiring very long runtimes in order to cover the relevant posterior mass. For synthetic likelihood-based approaches another issue is the large number of experimental measurements (16k), which significantly increases the variance of the total likelihood estimates and reduces acceptance rates even further. For these reasons we followed the approach of the authors in [24, 52], performing Maximum Likelihood Estimation (MLE) and comparing the predictive distribution with experimental data^1^.

The results can be seen in Fig. 4**b**. Since the data comes in the form of independent population snapshots we used independent mixtures of negative binomials as our synthetic model. Parameters obtained using FSP and our approach provide good agreement with the experimental data as reported in [24], whereas GSL and MBI often failed to capture the shape of the experimental distribution. These results show that synthetic models can be applied to obtain high-quality parameter estimates for real-life biochemical systems with comparable accuracy to FSP.

## 4 Discussion

We presented an approach for inference in stochastic gene regulatory networks relying on an approximation of the generally intractable CME by a family of synthetic models, fit to the original model via simulations. These synthetic models yield estimates of the model likelihood, which can be optimised to obtain maximum likelihood estimates for the true model parameters, or within an MCMC sampler for posterior inference and model selection.

We tested our method on a well-studied autoregulatory feedback loop and showed that it closely approximates the exact posterior in both the positive and the negative feedback regimes, recovering true parameters with significantly more accuracy than standard approaches such as Moment-Based Inference and Approximate Bayesian Computation. We then considered a more complex model, the genetic toggle switch, which is difficult to analyse using moment-based methods and the FSP, illustrating the flexibility of our approach and its ability to handle nontrivial models of real-life systems. Our findings show that distributional approximations beyond Gaussians can aid parameter identifiability, and that simulation-based methods can be effectively used in place of analytical approximations where the latter fail.

We finally demonstrated the effectiveness of our approach for analysing real-life data by testing it on the MAPK pathway in *S. Cerevisiae* in [52], obtaining parameter estimates rivalling those of the FSP in predictive accuracy. A significant advantage of our approach over the FSP is that its computational complexity grows only modestly in the number of species, whereas solving the FSP for models such as the genetic toggle switch is a difficult numerical problem. This allows synthetic models to be used for more realistic reaction networks that are challenging to analyse with the FSP, with a negligible trade-off in accuracy as exemplified by our results.

A major limitation of our method is that fitting a synthetic model to simulations introduces a variance in the approximate likelihood proportional to the number of experimentally observed datapoints. In order to obtain accurate estimates of the true likelihood, therefore, the number of simulations used to train the synthetic model needs to be increased in step with the sample size. For MLE estimation this does not significantly complicate things, but in an MCMC context this variance causes difficulties as it can heavily reduce acceptance rates. Another limitation of our method is that a MNB can only provide an accurate approximation for (transcript or protein) marginal distributions with a Fano factor greater than 1. This condition is met in the overwhelming majority of computational models and experimental studies of gene regulatory systems, but exceptions exist [53–55]. Incorporating a different parametric family of distributions with Fano factor smaller than 1 (e.g. hypergeometric) is in principle straightforward within the SM framework.

The Metropolis-Hastings sampler used in this paper is most suited for low-dimensional problems spaces as a random walk-based approach is not an efficient way to explore high-dimensional posteriors. MCMC sampling in high-dimensional spaces is often done using the Metropolis-adjusted Langevin Algorithm (MALA) or Hamiltonian Monte Carlo (HMC) [56], both of which require gradients of the posterior to direct the sampler towards high-probability regions. Approximating the gradient of the likelihood function using synthetic likelihoods is therefore a promising direction for future research.

The need to fit a synthetic model from scratch at every iteration of the MCMC procedure is the computational bottleneck of our method. Methods such as data subsampling [57, 58] and amortisation, e.g. using neural networks [35] could result in significant speedups and a reduced variance in the likelihood estimates for more complex problems.

An advantage of our approach over standard CME-based inference methods is that it can be readily applied to systems with extrinsic noise, simulated using the exact Extrande algorithm [59], and/or non-Markovian systems such as those considered in [40, 41, 60, 61]. While such models are difficult to analyse mathematically, requiring various extensions to the CME formalism, the presence of efficient and exact versions of the SSA for these systems allows most simulator-based inference methods to work without any modification. We hope that our work, as well as the ideas contained within, provides a useful stepping stone that will enable researchers to analyse and use these models more efficiently in the future.

## Materials and Methods

### Gaussian Synthetic Likelihoods

We model the distribution of observed molecule numbers, jointly at all time points, as a multivariate Gaussian whose mean and covariance we estimate from simulations obtained using the SSA.

### Moment-Based Inference

The moments of the CME can be computed at any point in time from its associated moment equations. For linear systems with mass-action kinetics, such as the MAPK pathway example, these can be solved directly, while for the autoregulatory feedback loop and the genetic feedback loop we used the Linear Mapping Approximation [49] to obtain a solvable set of equations.

Experimentally observed moments form a stochastic estimate of the true moments of the system; for large sample sizes, the Central Limit Theorem ensures that these sample moments will be approximately normally distributed. Following the approach in [17, 23] we thus model the first and second (uncentered) sample moments over observed molecule numbers using a multivariate Gaussian. The means and covariances of the sample moments can be expressed in terms of the analytical moments of the system, and we assume that measurements at different timepoints are independent (see SI for details). This results in a Gaussian likelihood on the moment level that can be used for inference.

### Approximate Bayesian Computation

We use the first and second-order moments over species numbers at each time point as summary statistics. Fixing a tolerance *ϵ* we repeatedly sample parameters from the prior and compare the simulator output **x** with the observed data **x**_obs_. Namely we accept parameters for which the sum of the squared relative errors in the first and second moments is less than *ϵ* and iterate until a prespecified number of acceptances is reached. To improve sample efficiency we decrease *ϵ* over multiple rounds following [28], using a Gaussian proposal prior estimated from the results of the previous round to guide sampling. Regression adjustment [62] did not yield measurable improvements in our experiments.

## Code Availability

Code implementing synthetic models as well as the experiments in this paper is available at https://github.com/kaandocal/synmod.

## Acknowledgments

K. Ö. was supported in part by the EPSRC Centre for Doctoral Training in Data Science, funded by the UK Engineering and Physical Sciences Research Council (grant EP/L016427/1) and the University of Edinburgh. R. G. and G. S. acknowledge support from the Leverhulme Trust (RPG-2018-423). K. Ö. and R. G. would like to thank Guillaume Lieb for constructive discussions of the MAPK pathway model.

## Footnotes

1 We emphasise that the predictive distributions are obtained by running the model at the estimated parameters and are *not* mixtures of negative binomials (for SM) or Gaussians (for GSL).

## Notes

### Competing Interest Statement

The authors have declared no competing interest.

