## Supplementary material for "Inference and Uncertainty Quantification of Stochastic Gene Expression via Synthetic Models": SI


#### 1 Fitting Synthetic Models

In order to fit a mixture of negative binomials to data we use the Expectation-Maximisation algorithm described in [1, 2]. Let $x_1, \dots, x_N$ be a set of independently sampled observations from an unknown mixture of negative binomials, and let $z_j$ denote the (latent) mixture component each $x_j$ was sampled from. The parameters of the mixture components are given by $r_i$, $p_i$ and weights $w_i$ for $i = 1, \dots, m$.

Expectation-Maximisation is an iterative algorithm that refines the estimates of these parameters across rounds using the following formulae:

$$
\begin{aligned}
z_{ij}^{(k)} &:= p(z_j = i \,|\, x_j) \propto w_i^{(k)} NB(x_j;\, r_i^{(k)}, p_i^{(k)}), \\
\delta_{ij}^{(k)} &:= r_i^{(k)} \left( \Psi(r_i^{(k)} + x_j) - \Psi(r_i^{(k)}) \right), \\
\beta_i^{(k)} &:= 1 - \frac{1}{1 - p_i^{(k)}} - \frac{1}{\log p_i^{(k)}}.
\end{aligned}
$$

Here the superscript indicates the current round. The updated parameters are given by

$$
w_i^{(k+1)} = \sum_j \tau_{ij}^{(k)},
$$

$$p_i^{(k+1)} = \frac{\beta_i^{(k)} \sum_j \tau_{ij}^{(k)} \delta_{ij}^{(k)}}{\sum_j \tau_{ij}^{(k)} (x_j - (1 - \beta_i^{(k)}) \delta_{ij}^{(k)})},$$

$$r_i^{(k+1)} = -\frac{1}{p_i^{(k+1)}} \frac{\sum_j \tau_{ij}^{(k)} \delta_{ij}^{(k)}}{\sum_j \tau_{ij}^{(k)}}.$$

This iteration step is repeated until convergence is achieved, which can be seen as either the parameters or the log-likelihood stop changing significantly.

The above procedure can be combined with the Baum-Welch algorithm to fit a HMM with negative binomial emission distributions. The transition rates and initial distribution are fit as usual, and the emission distributions are updated using the above procedure for $p_i$ and $r_i$ with $z_i j = p(z_j = i \,|\, x_j)$ computed with the forward-backward algorithm.

### 2 Numerical Experiments

We used Julia's `Catalyst.jl` [3] for defining reaction networks, `DifferentialEquations.jl` [4] for stochastic simulations and solving ODEs, `BlackBoxOptim.jl` for optimisation, `GpABC.jl` for SMC-ABC and `MomentClosure.jl` [5] for creating moment equations. Code for all experiments is available on GitHub.

MCMC experiments were initialised by choosing random parameters in the prior and optimising for 1000 steps to find relevant regions of the posterior. Optimisations were performed using the default Adaptive Differential Evolution method implemented in `BlackBoxOptim.jl`. We used the standard Metropolis-Hastings algorithm with a symmetric Gaussian transition kernel.

All experiments were performed on an Intel Xeon Silver 2.20Ghz machine with 20 cores. Unless otherwise noted SSA simulations were initialised with 0 mRNA/protein and one copy of each gene in the unbound state.

##### Autoregulatory Feedback Loop

With the FSP we used a truncation to $\leq 150$ proteins for the positive feedback loop and $\leq 200$ proteins for the negative feedback loop. With SM we used $5$ mixture components and 5k samples per likelihood evaluation, with GSL we used 3k samples.

We ran the Metropolis-Hastings algorithm for 50k iterations with each method after a burn-in of 5k samples. The transition kernel was chosen to be diagonal Gaussian with standard deviation a fixed fraction of the prior standard deviation in each dimension. For FSP and SM the fraction was 5% in each dimension, except for $\rho_b$ for which it was 1%. For GSL the fraction was 10% in each dimension, and for MBI 15%.

Each MCMC experiment was done in a few hours, with moment-based inference taking several minutes. Maximum-Likelihood Estimates were taken by picking the posterior sample with the highest computed likelihood. For ABC we used mean and standard deviation of protein numbers at each time point as summary statistics, with discrepancy measure the square root of the sum of squared relative errors. The tolerance schedule for ABC was $\epsilon = 1.5, 1.25, 1.0, 0.75$.

##### Genetic Toggle Switch

With SM we used $4$ mixture components, and both SM and GSL were fit using 10k samples per likelihood evaluation. We ran the Metropolis-Hastings algorithm for 500k iterations (800k for GSL due to the lower acceptance rate) after a burn-in of 5k. The transition kernel was chosen to be diagonal Gaussian with standard deviation a $1\%$ of the prior standard deviation in each dimension. Maximum-Likelihood Estimates were taken by picking the posterior sample with the highest computed likelihood.

For ABC we used means, variances and covariance of the numbers of protein A and B at each time point as summary statistics, with discrepancy measure the square root of the sum of squared relative errors. The tolerance schedule for ABC was $\epsilon = 20, 15, 10, 7.5$.

**MAPK Pathway Model**

Experimental data for this system were obtained from the authors of [6]. We used the wild-type strain of the STL1 subjected a 0.4M NaCl osmotic shock the kinase concentration was modelled using the deterministic $\mathrm{Hog1p}^*(t)$ function described in the paper.

For the FSP we used a truncation to $\leq 150$ mRNA molecules to compute marginal mRNA distributions at each time point. For SM we used $10$k simulated samples and $4$ mixture components, fitting mixtures independently at each time point due to the fact that we have population snapshot data. For GSL we used $10$k samples and fit independent Gaussians at each time point for the same reason. Due to the cheapness of the FSP in this example samples were obtained via the FSP rather than the Gillespie Algorithm. For MBI we used the first two moments of the observed distribution at each time point.

Each parameter was restricted to $[0.001, 1000]$, except $b21$ which was constrained to $[-1e5, 1e5]$, negative values representing kinase-induced inhibition. Optimisation was performed in the Julia package `BlackBoxOptim.jl`, using the default algorithm (Adaptive Differential Evolution method) with a maximum of 25k steps. We report the best of three runs for each method, each of which took a few hours except MBI (less than an hour).

**a** **Estimated Parameters**

| | GT | FSP | SM | GSL | MBI |
|---|---|---|---|---|---|
| $\sigma_u$ | 0.1 | 0.047 | 0.087 | 0.087 | 0.0978 |
| $\sigma_b$ | 0.001 | 0.0012 | 0.0012 | 0.0016 | 0.0019 |
| $\rho_u$ | 13.0 | 13.8 | 13.0 | 13.4 | 50.0 |
| $\rho_b$ | 0 | $< 0.001$ | 0.001 | 0.03 | 0.2 |
| $b$ | 3 | 2.75 | 3.0 | 2.9 | 0.81 |

**b** **Mean TV Distances**

| SM | GSL | MBI | ABC |
|---|---|---|---|
| 0.124 | 0.244 | 0.201 | 0.206 |

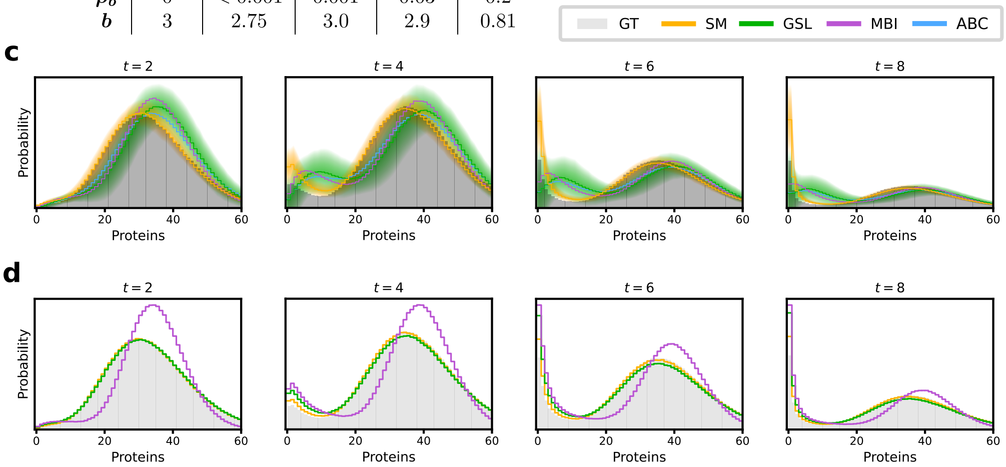


**Figure 1:** Inference results for the negative autoregulatory feedback loop. **a**: Ground truth parameters and maximum likelihood estimates obtained using the Finite State Projection, Synthetic models, Gaussian synthetic likelihoods, and moment-based inference. **b**: Mean total variation distance between the predicted distribution and the true distribution for the posteriors obtained using different methods, averaged over the time points shown. Results were obtained by taking 100 parameter sets from each posterior. **c**: Posterior predictive protein number distributions at different time points. Solid lines denote the posterior mean predictive distribution and the shaded areas show one standard deviation. **d**: Predictive protein distributions for the maximum likelihood parameters inferred using SM, GSL and MBI.

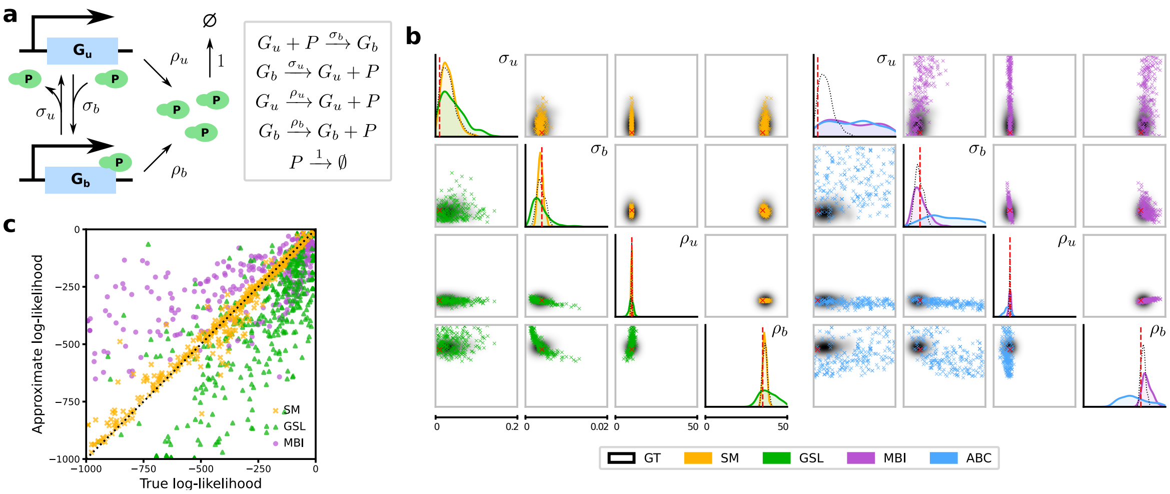


**Figure 2:** Comparison of Synthetic Models with standard methods for the case of an autoregulatory genetic positive feedback loop. **a**: Illustration of the reaction scheme. Note that protein is here produced in bursts of size 1 – this is a special case of the bursty model in Fig. 2a of the main text; for a derivation see [7] **b**: Posteriors obtained using four different inference methods, with the FSP solution indicated in black. The ranges plotted coincide with the prior ranges. **c**: Comparison of true and approximate log-likelihoods. Parameter values were sampled from the prior and the true log-likelihoods were computed using the FSP. The true parameter values are given in Fig. 3. The input data consists of protein numbers from 25 SSA trajectories measured at times $t = 4, 8, 12, 16$.

**a** **Estimated Parameters**

| | GT | FSP | SM | GSL | MBI |
|---|---|---|---|---|---|
| $\sigma_u$ | 0.01 | 0.023 | 0.018 | 0.016 | < 0.001 |
| $\sigma_b$ | 0.004 | 0.0034 | 0.0035 | 0.004 | 0.0023 |
| $\rho_u$ | 10 | 9.8 | 9.8 | 9.3 | 10.2 |
| $\rho_b$ | 35 | 36.3 | 35.3 | 34.4 | 37.3 |

**b** **Mean TV Distances**

| SM | GSL | MBI | ABC |
|---|---|---|---|
| 0.094 | 0.197 | 0.174 | 0.340 |

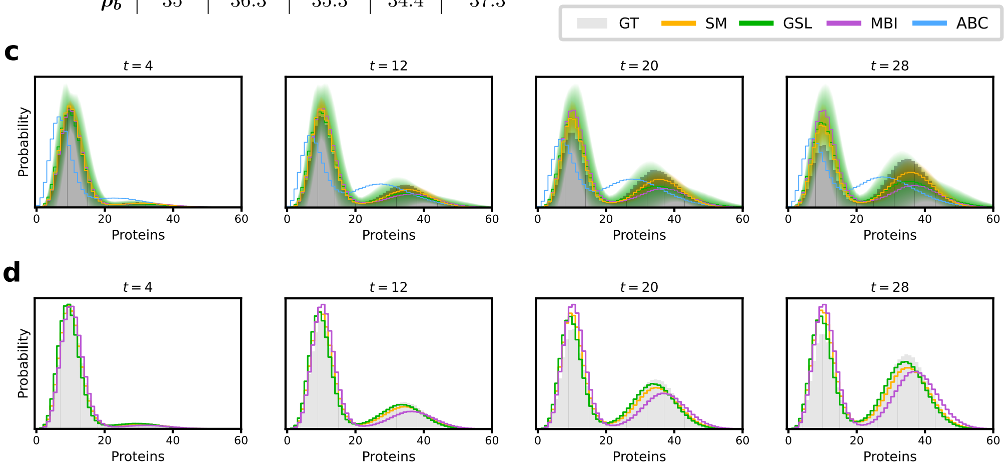


**Figure 3:** Inference results for the positive autoregulatory feedback loop, cf. Fig. 1. **a**: Ground truth parameters and maximum likelihood estimates. **b**: Mean total variation distance between the predicted distribution and the true distribution for the posteriors, averaged over the time points shown. Results were obtained by taking 100 parameter sets from each posterior. **c**: Posterior predictive protein number distributions at different time points. **d**: Predictive protein distributions for the maximum likelihood parameters inferred using SM, GSL and MBI.

**a**

**Estimated Parameters**

| | | GT | SM | GSL | MBI | Prior Bounds |
|---|---|---|---|---|---|---|
| Gene A | $\sigma_u$ | 6 | 4.6 | 10.9 | 0.2 | [0, 20] |
| | $\sigma_b$ | 12 | 9.0 | 19.6 | 0.1 | [0, 40] |
| | $\rho_p$ | 15 | 12.9 | 8.6 | 19.1 | [0, 50] |
| | $\rho_p$ | 30 | 34.5 | 50.7 | 66.0 | [0,100] |
| Gene B | $\sigma_u$ | 7 | 11.4 | 13.7 | 6.1 | [0, 25] |
| | $\sigma_b$ | 1.5 | 2.5 | 2.8 | 0.7 | [0, 8] |
| | $\rho_p$ | 1.7 | 1.8 | 1.5 | 2.3 | [0, 10] |
| | $\rho_p$ | 60 | 53.4 | 64.0 | 104 | [0, 200] |

**Fixed Parameters**

| | | GT | | | GT |
|---|---|---|---|---|---|
| Gene A | $D_m$ | 12 | Gene B | $D_m$ | 16 |
| | $\delta_m$ | 3 | | $\delta_m$ | 1 |
| | $D_p$ | 4 | | $D_p$ | 4 |
| | $\delta_p$ | 2 | | $\delta_p$ | 3 |

**b**

**Mean TV Distances**

| | SM | GSL | MBI | ABC |
|---|---|---|---|---|
| $P_A$ | 0.044 | 0.080 | 0.167 | 0.139 |
| $P_B$ | 0.040 | 0.051 | 0.218 | 0.114 |

**c**

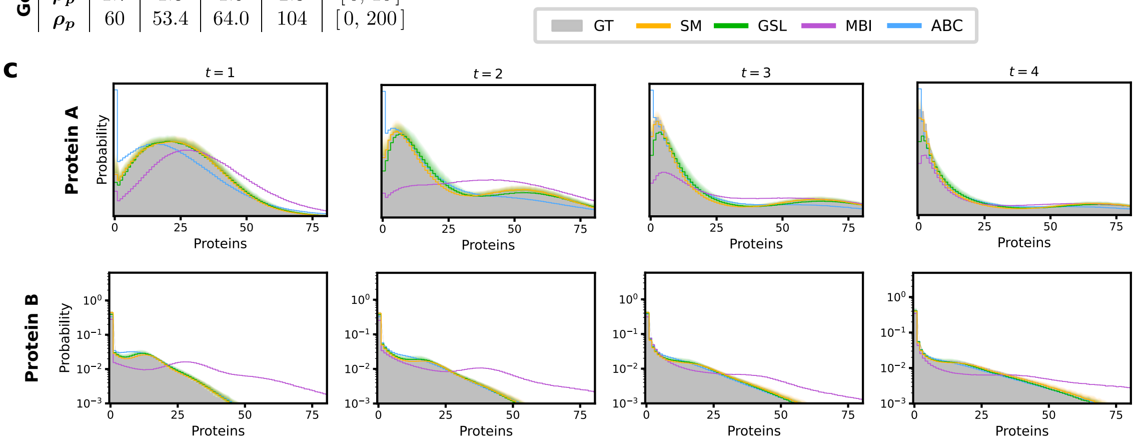


**Figure 4:** Inference results for the toggle switch. **a**: Ground truth parameters, maximum likelihood estimates and prior bounds for the free parameters. **b**: Mean total variation distance between the predicted protein distributions and the true distribution for the posteriors obtained using different methods, averaged over the time points shown. Results were obtained by taking 100 parameter sets from each posterior. **c**: Posterior predictive protein number distributions at different time points. Solid lines denote the posterior mean predictive distribution and the shaded areas show one standard deviation.

| | FSP | SM | GSL | MBI |
|---|---|---|---|---|
| $\boldsymbol{k_{12}}$ | 3.7 | 0.14 | 0.01 | 0.017 |
| $\boldsymbol{a_{21}}$ | 695 | 768 | 672 | 168 |
| $\boldsymbol{b_{21}}$ | -3539 | -2904 | -2490 | -1267 |
| $\boldsymbol{k_{23}}$ | 0.0078 | 0.033 | 0.0018 | 0.97 |
| $\boldsymbol{k_{32}}$ | 4.75 | 0.019 | 0.01 | 49.4 |
| $\boldsymbol{k_{34}}$ | 667 | 0.013 | 0.076 | 0.014 |
| $\boldsymbol{k_{43}}$ | 2.2 | 0.018 | 86.5 | 0.91 |
| $\boldsymbol{\rho_1}$ | 0.0023 | 0.0013 | 0.0023 | 0.0019 |
| $\boldsymbol{\rho_2}$ | 0.016 | 0.032 | 0.11 | 0.040 |
| $\boldsymbol{\rho_3}$ | 125.6 | 0.002 | 0.49 | 0.13 |
| $\boldsymbol{\rho_4}$ | 0.0017 | 0.44 | 0.036 | 625 |
| $\boldsymbol{\delta}$ | 0.0054 | 0.0052 | 0.0050 | 0.0061 |

**Table 1:** Estimated parameters for the MAPK pathway model.
